# Common postzygotic mutational signature in multiple healthy adult tissues related to embryonic hypoxia

**DOI:** 10.1101/2020.02.17.952473

**Authors:** Yaqiang Hong, Dake Zhang, Xiangtian Zhou, Aili Chen, Amir Abliz, Jian Bai, Liang Wang, Qingtao Hu, Kenan Gong, Xiaonan Guan, Mengfei Liu, Xinchang Zheng, Shujuan Lai, Hongzhu Qu, Fuxin Zhao, Shuang Hao, Zhen Wu, Hong Cai, Shaoyan Hu, Yue Ma, Junting Zhang, Yang Ke, Qianfei Wang, Wei Chen, Changqing Zeng

**Affiliations:** Key Laboratory of Genomic and Precision Medicine of Chinese Academy of Sciences, Beijing Institute of Genomics, Beijing, China; Beijing Advanced Innovation Center for Biomedical Engineering, School of Biological Science and Medical Engineering, Beihang University, Beijing, China; Tsinghua-Peking Center for Life Sciences, School of Life Sciences, Tsinghua University, Beijing, China; School of Optometry and Ophthalmology and Eye Hospital, Wenzhou Medical University, Wenzhou, Zhejiang, China; State Key Laboratory Cultivation Base and Key Laboratory of Vision Science, Ministry of Health P. R. China and Zhejiang Provincial Key Laboratory of Ophthalmology and Optometry, Wenzhou, Zhejiang, China; Key Laboratory of Carcinogenesis and Translational Research (Ministry of Education), Laboratory of Genetics, Peking University Cancer Hospital & Institute, Beijing, China; Skull Base and Brainstem Tumor Division, Department of Neurosurgery, Beijing Tian Tan Hospital, Capital Medical University, Beijing, China; China National Clinical Research Center for Neurological Diseases, NCRC-ND, Beijing, China; Key Laboratory of Genome Sciences and Information of Chinese Academy of Sciences, Beijing Institute of Genomics, Beijing, China; Pediatrics, Hematology and Oncology, Children’s Hospital of Soochow University, Soochow, Suzhou, China; Institute of Biophysics, Chinese Academy of Sciences, Beijing, China; Collaborative Innovation Center for Genetics and Development, Shanghai, China; University of Chinese Academy of Sciences, Beijing, China

## Abstract

Postzygotic mutations are acquired in all of the normal tissues throughout an individual’s lifetime and hold clues for identifying mutagenesis causing factors. The process and underlying mechanism of postzygotic mutations in normal tissues is still poorly understood. In this study, we investigated postzygotic mutation spectra in healthy individuals by optimized ultra-deep exome sequencing of time series samples from the same volunteer and samples from different individuals. In cells of blood, sperm, and muscle, we resolved three common types of mutational signature. Two of them are known to represent clock-like mutational processes, and their proportions in mutation profiles associated with polymorphisms of epigenetic regulation genes, suggesting the contribution of personal genetic backgrounds to underlying biological process. Notably, the third signature, characterized by C>T transitions at GpCpN sites, tends to be a feature of diverse normal tissues. Mutations of this type were likely to occur early in embryo development even before the tissue differentiation, as indicated by their relatively high allele frequencies, sharing variants between multiple tissues, and lacking of age-related accumulation. Almost all tumors shown in public datasets did not have this signature detected except for 19.6% of clear cell renal cell carcinoma samples, which featured by activation of the hypoxia-induced signaling pathway. Moreover, in vitro activation of HIF signaling pathway successfully introduced the corresponding mutation profile of this signature in a culture-expanded human embryonic stem cell line. Therefore, embryonic hypoxia may explain this novel signature across multiple normal tissues. Our study suggest hypoxic conditions in the early stage of embryo development may be a crucial factor for the C>T transitions at GpCpN sites and individual genetic background also related to shaping human postzygotic mutation profiles.

## Introduction

After fertilization, most genomic mutations typically occur as a result of replication errors, DNA structure instability, as well as other endogenous and exogenous sources, resulting in the genotypic and phenotypic heterogeneity of all types of cells in the body [1-3]. In particular, mutations can be triggered by distinct environmental factors, producing characteristic patterns. The accumulation of somatic mutations is believed to chronicle the exposures, toxicity, regeneration and clonal structure of the progresses from health to disease [4-6]. Thus, the roles of somatic mutations in pathogenesis have been widely explored [7, 8]. Moreover, in recent years, multiple cell clones with distinct genotypes, referred to as somatic mosaicism by lineage expansion in healthy tissues, have drawn attention to the factors underlying certain disorders [9, 10].

Tissue-specific processes or particular microenvironmental changes leave unique imprints in genomes [9, 11]. With the advent of next-generation sequencing (NGS), characteristics of multiple mutagenic processes have been revealed for the first time in tumors of various origins [7, 11-13]. For instance, smoking results mainly in C>A transitions in lung cancers, while ultraviolet (UV) radiation leaves a footprint involving CC>TT dinucleotide substitutions in skin cancers [7, 14]. A recent investigation showed distinct mutational spectra in cultured adult stem cells (ASCs) of liver in comparison with those originating from the colon and small intestine [9]. Moreover, mutation spectra are influenced by the genetic background of individuals. For example, breast cancer patients having *BRCA1* or *BRCA2* germline mutations showed a specific mutational signature in tumor genomes compared with patients carrying *BRCA* wild types [11]. The confounding of different mutagenesis-related factors by the genetic background means that mutation accumulation patterns differ among tissues and individuals.

Two mutational signatures (Signature 1 and Signature 5 in COSMIC) related to the deamination of methylated cytosines have shown a feature of accumulation with age in a broad range of cell types. However, the accumulation process does not seem to maintain a steady pace. Specifically, the mutation rate per cell division varies during development, undergoing diverse biological changes prenatally, and in childhood and adulthood [1]. *De novo* mutations in offspring increase with paternal age, and the accumulation rate in gonads was estimated to be ∼2 mutations per year [15]. More than twofold differences in variation have been observed between families, possibly influenced by germline methylation [1]. Hence, factors that influence the mutagenic processes may differ due to various developmental demands, such as the activities of stem cells in tissue repair, exposure to environmental factors, and tissue-specific functions [7, 9]. In addition, the changes in mutational profile in cultured cells also reflect the genetic drift that occurs during clonal expansion of the cell population carrying multiple pre-existing mutations [16].

The large majority of knowledge on somatic mutation has been obtained from genomic analyses of cancer or noncancer diseases, animal models, and cultured cells. However, despite the importance of analyzing the generation and subsequent effects of somatic mutations in normal tissues, studies on their mutation profiles are limited due to not only difficulties in obtaining appropriate tissues from healthy individuals, but also the scarcity of cells carrying mutations [17-19]. Although great effort has been made on analyzing somatic mutation profile on various tissue including skin, liver, esophagus, and colon, our knowledge of the mutation spectrum and its dynamic nature in healthy individuals remains inadequate [6, 10, 20, 21].

To obtain the somatic mutation spectrum in healthy individuals, in this study we first conducted optimized ultra-deep exome sequencing (∼800×) of blood samples in five trio families. From deep sequencing for time points samples of blood, muscle and sperm in one subject, followed by comparison with results of another 50 samples, we identified a mutational signature characterized by C>T transition at GpCpN, specific to normal tissues. Further association analysis suggested that certain SNPs residing in epigenetic regulators may explain the individual-specific proportions of C>T at GpCpN in the population. An *in vitro* experiment and somatic mutation data from cancer genome research further showed that hypoxia is a trigger for mutagenesis.

## Results

### Postzygotic mutations in normal blood and sperm cells revealed by ultra-deep exome sequencing

We adopted ultra-deep exome sequencing (>800× coverage) to identify genomic mutations, with the benefits of increased sensitivity and accuracy due to multiple steps of optimization (S1 Fig, Methods, S1 Appendix). First, we analyzed five specimens from the volunteer M0038 annually for 4 years, including 2 blood and 3 sperm samples (S1 Table). In both tissues, one *de novo* mutation was identified and its variant allele fraction (VAF) reached 0.4 +/- 0.02 in all samples. Overall, 36 cross-tissue mutations, with VAF of 0.002 to 0.434, were shared by at least one blood and one sperm sample (Fig 1A, S2 Fig, and S2 Table). They were likely to occur before tissue differentiation, considering the previous speculation that some cells may contribute to multiple tissues at the early stages of embryo development [22]. For tissue-specific mutations, four common postzygotic mutations were detected in all of the whole-blood samples, and 16 common mutations were seen in all three sperm samples (Fig 1A-B). Especially, VAFs for these mutations were all consistent across the samples.

**Fig. 1.**
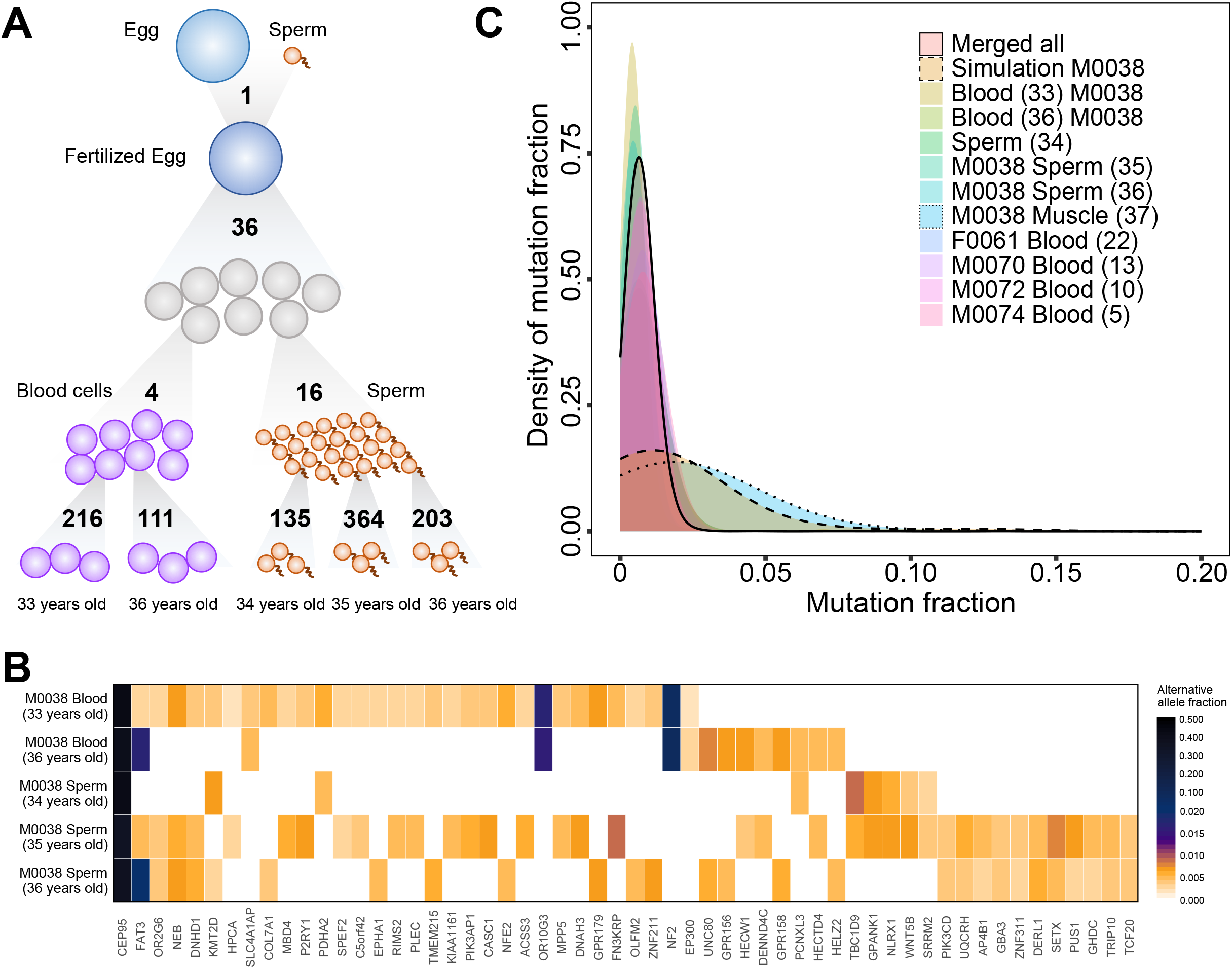
Postzygotic mutation profiling. (A) Schematic diagram depicting mutation accumulation among the time-point samples from individual M0038. Each tested sample carries a couple of hundred mutations as private events (numbers at the bottom). One *de novo* mutation occurred before fertilization (top). Thirty-six mutations were shared by at least one sperm sample and one blood sample. In contrast with the 16 shared common mutations found in sperm samples but not in blood, and only four blood specific mutations were shared by both ages in blood. (B) Shared mutated genes in different samples of M0038. Among the 57 mutations revealed in at least two samples besides the *de novo* mutation (*CEP95*), only one mutation (*NF2*, 0.09) was identified to have an allele fraction greater than 0.05. The scaled color represents the allele fraction of mutations. (C) Density plot of mutation fraction distribution. A significantly higher density was shown in the low-fraction region of all samples (dashed pink area) and there were more mutations with low allele fractions than the mutations obtained by muscle (light blue area with dotted line) and simulation (beige area with dashed line).

We further compared postzygotic mutation profiles in five trio families with this approach (including M0038). Overall, 3,266 postzygotic mutations and 4 *de novo* mutations (Methods) in blood samples from children in the five trios were detected with VAF ranging from 0.002 to 0.528 (S3 Fig, S3 Table), and the validation rate was above 85% using multiple methods (Methods and S2 Appendix). Approximately 90% of the variations in all individuals had allele fractions of less than 0.020, indicating that only a small subset of cells carried the mutations. Based on the assumption that the mutation rate during cell divisions is stable, *in silico* simulation [23, 24] showed a range of allele fractions from 0.050 to 0.200 (beige area in Fig 1C, S4 Fig, and Methods), but our observations demonstrated a significant difference from the simulation (p < 2.2×10^−16^, Kolmogorov–Smirnov test). This suggested the mutation rate may be unequal during cell divisions, consistent with the findings in previous studies on embryonic development [1, 25]. Our sample size limited the power to detect the relationship between amounts of postzygotic mutations and age, the former of which varied from 124 to 813 (S1 Table). Besides, these mutations had low recurrence rates. On average, only 2.7 (range 0–7) mutations were shared by two individuals (S2 Fig), and none was found in more than two individuals (S3 Table).

### Significant enrichment of GpCpN and NpCpG postzygotic mutations in normal tissues

We summarized the trinucleotide composition of all 96 substitution types in each individual according to their position and two neighboring bases. As demonstrated in Fig 2A, C>T transitions were enriched in all individuals, of which more than 90% were in GpCpN or NpCpG sites in both blood and sperm cells (S5 Fig). For these two trinucleotide contexts, only individual M0038 had more GpCpN than NpCpG mutations in all types of samples, whereas the other individuals had more NpCpG mutations. This suggests the existence of distinct mutational processes among individuals.

**Fig. 2.**
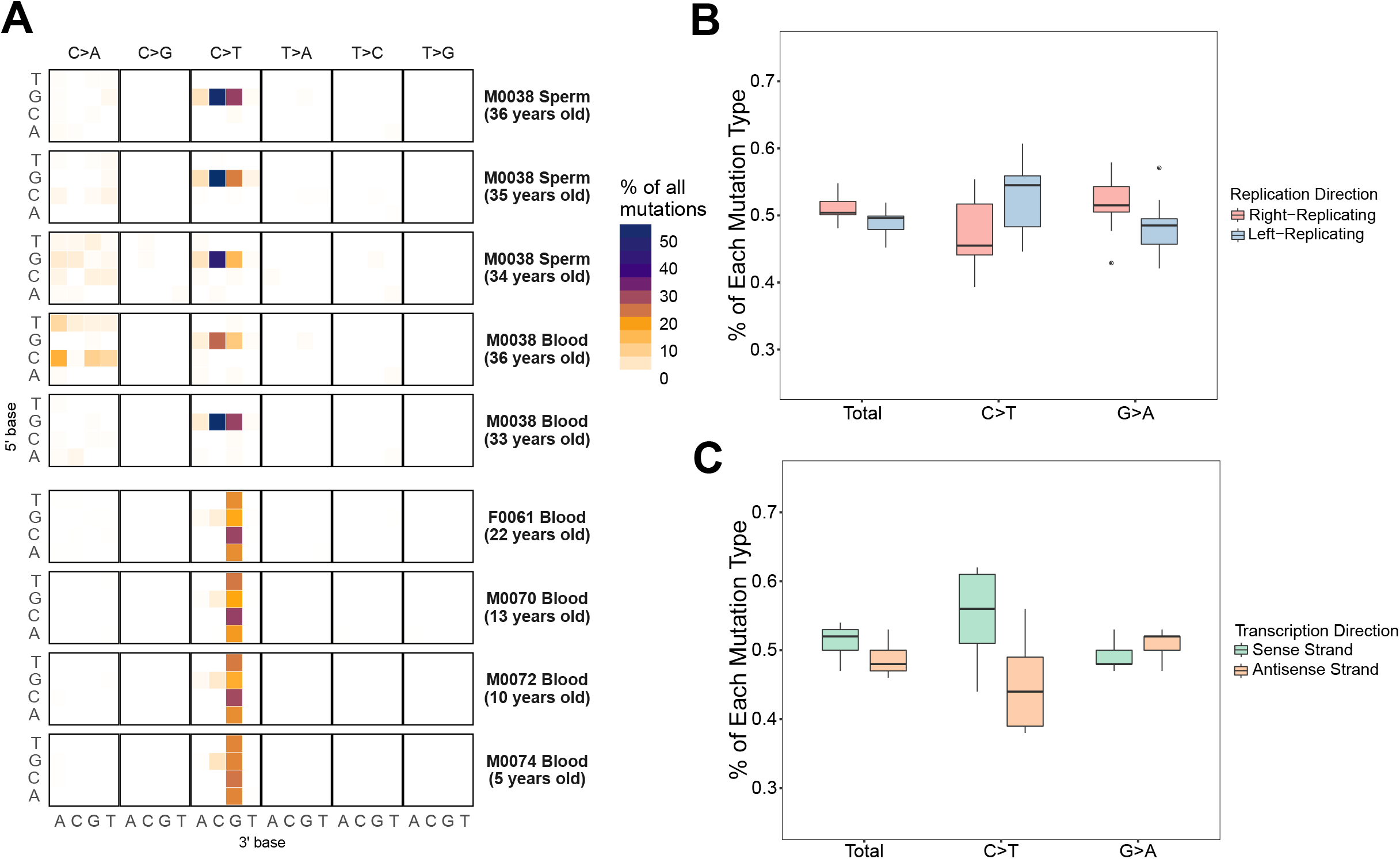
Patterns of postzygotic mutations in healthy individuals. (A) Heat map of the rates of each mutation type. Significant enrichment of C>T transitions, especially at NpCpG and GpCpN sites, was exhibited at each of the 96 mutated trinucleotides in all individuals. Similar patterns were shown among various types of sample from individual M0038. C>A transversions with no preferred context were also detected in normal cells. (B) Strand asymmetry of C>T transitions by analysis of the replication direction. C>T transitions were more likely to occur in the left-replication regions and G>A were enriched in the right-replication regions, suggesting mutational strand asymmetry due to replication. (C) The transcription asymmetry of C>T transitions. C>T was also more likely to occur in regions with the sense strand as the encoded strand, whereas G>A exhibited high enrichment in the opposite regions.

C>T transitions at NpCpG sites commonly originate from age-related spontaneous deamination of methylated cytosine to thymine [2, 7]. Nevertheless, time-point samples for the individual M0038 did not show the time-related feature of C>T transitions at NpCpG, with their proportions varying from 13% to 31% (S6 Fig). Among other four individuals, their counts slightly increased with age but without any significance (e.g., 94% in the youngest individual, 5-year-old M0074, and 95% in 22-year-old F0061; S6 Fig). Meanwhile, the proportions of C>T at GpCpN were consistent across all samples, with the highest rate of 29% and the lowest of 23% (S6 Fig).

Moreover, these mutations also demonstrated reported mutational strand asymmetries caused by replication and transcription (Methods). For the C>T or G>A transition at GpCpN and NpCpG, the C>T transitions were likely to occur in the left-replication regions of the genome during DNA replication, whereas more G>A transitions occurred in the right-replication regions (Fig 2B and S7 Fig). The genomic regions that encoded genes on the reference strand exhibited a high density of C>T transitions, and the regions that encoded genes on the complementary strand exhibited a high density of G>A transitions (Fig 2C and S8 Fig). Additionally, sperm and blood cells from M0038 exhibited no significant difference in the patterns of postzygotic mutations in the 96 mutation contexts and the mutational strand asymmetries (Fig 2A and S6 Fig – S8 Fig). Both tissues had higher levels of C>T transition at GpCpN than at NpCpG, indicating the same mutational processes. In brief, consistent VAF of mutations in samples of time series, similar proportions of C>T at GpCpN across samples from different individuals, and evidence of mutational strand asymmetries gave support to the reliability of mutation profiles we observed.

### A mutational signature characterized by C>T at GpCpN commonly occurred in normal blood cells

To explore the whether these mutation patterns were also represented in other normal tissues, we collected deep exome sequencing datasets (>200×) for one muscle sample from individual M0038 and normal blood cell samples paired to tumor samples in three types of tumors, including esophageal squamous cell carcinoma (ESCC), acute myelocytic leukemia (AML), and chordoma (Fig 3A). In addition, targeted sequencing data for normal skin and single cell sequencing data for neurons were also analyzed (Fig 3A). The most significant mutation feature observed in normal blood cells in these tumor studies was the enrichment of C>T transitions at GpCpN sites, especially the GpCpC trinucleotide, consistent with above profiles in healthy individuals. This kind of enrichment was also observed in the solid tissues, especially in neuron and muscle. Besides, all paired tumor samples did not have this feature detected in their mutation profiles. These strongly suggests that C>T at GpCpN sites commonly occurs in normal cells.

**Fig. 3.**
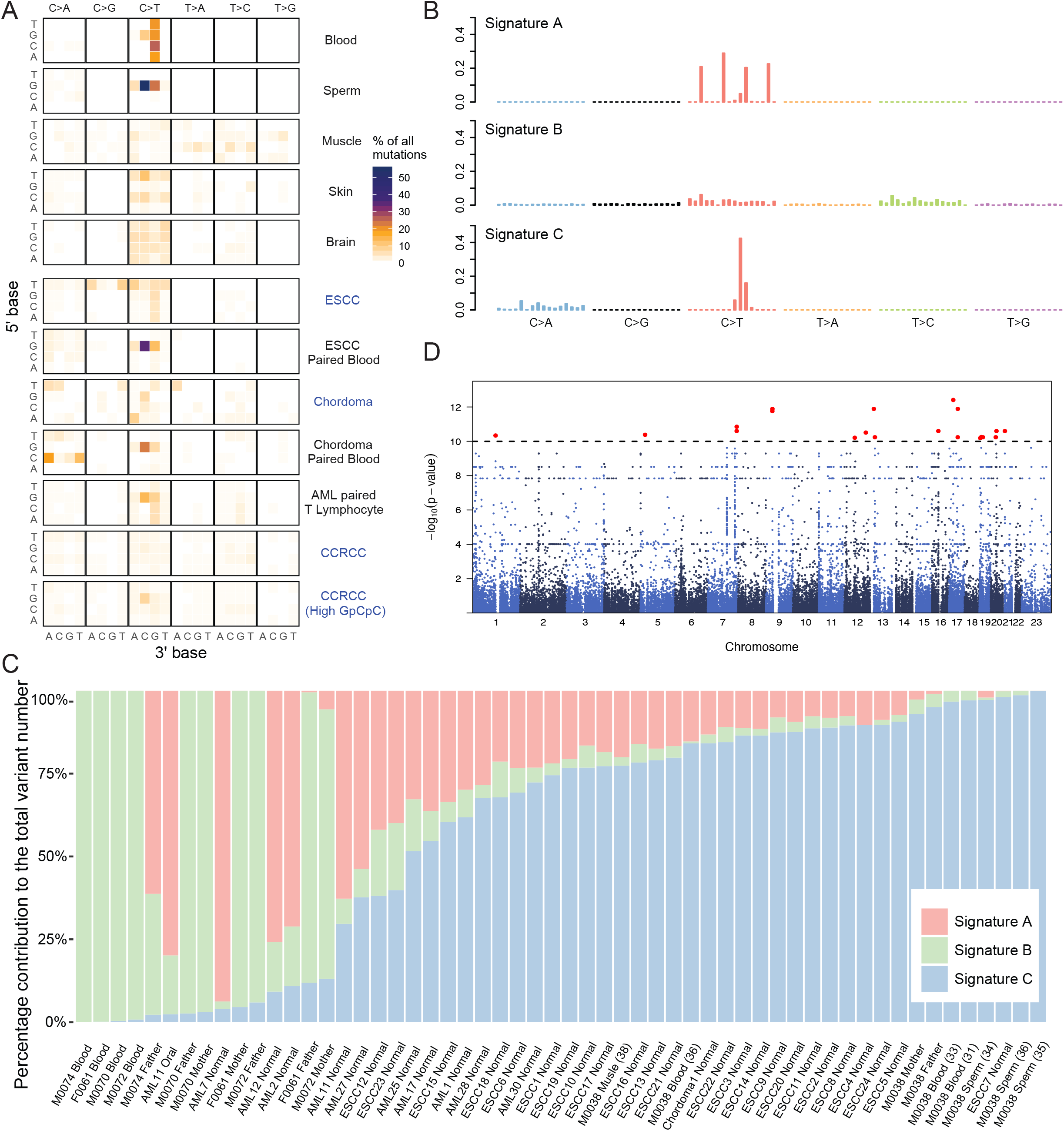
C>T at GpCpN sites in normal and tumor cells. (A) Heat map of mutation proportions illustrates the enrichment of C>T at GpCpN in all types of normal cell (black) in both healthy subjects (upper) and patients (lower). With the exception of enrichment of C>T at GpCpC in a portion of CCRCC, no specific mutation preference was identified in the trinucleotide contexts of various cancer cells (blue, note: data of AML sequencing are presented only for normal T lymphocytes, but not leukemic cells). However, distinct enrichment of C>T at GpCpN was demonstrated in all normal cells from both healthy individuals (upper 4) and paired blood samples in tumor patients (black in lower part). The mutation data presented from top to bottom are derived from the following sources: six blood samples of healthy individuals in this study; three sperm samples of subject M0038; one muscle sample of subject M0038 (>200×, VAF = 0.021±0.015); targeted sequencing (>500×, VAF = 0.042±0.048) of 74 genes in 234 skin samples from four individuals as reported by Martincorena *et al*. [5]; whole-genome sequencing of 36 single neurons of three individuals as reported by Lodato *et al*. [26]; in-house exome sequencing of 23 ESCC tumors and paired blood samples (both >200×, VAF = 0.017±0.009 in paired blood samples, VAF = 0.175±0.136 in tumors); in-house exome sequencing of two chordoma tumors and paired blood samples (both >250×, VAF = 0.017±0.005 in paired blood samples, VAF = 0.295±0.200 in tumors); in-house exome sequencing (>200×, VAF = 0.019±0.008) of 11 samples of normal T lymphocytes that were paired with AML cells (data not shown); exome sequencing of 295 CCRCC from TCGA; and those CCRCC with a high rate of C>T transition at GpCpC sites among the 295 samples. (B) A mutational signature revealed by non-negative matrix factorization in all 56 normal cell samples. Signature A involves spontaneous deamination of 5mC at NpCpG, Signature B features C>T and T>C transitions and Signature C features a mutational type characterized as C>T at GpCpN sites. (C) Varying proportions of the three signatures in 56 normal samples. Signature C contributed to at least 10% of the mutations in all samples and was the major mutational type in ∼30 samples. (D) With the proportion of Signature B as the quantitative value, 125 SNPs located in 54 genes correlated with the proportion of signatures by whole-exome association analysis (permutation test, p < 1×10^−10^).

Next, we merged deep exome sequencing datasets of all 56 normal tissue samples (Fig 3A, Methods), and resolved Three mutational signatures, A, B and C (Fig 3B). Signature A is known to be associated with the spontaneous deamination of methylated cytosine to thymine at NpCpG [7, 11, 27], and Signature B is known to be clock-like that the number of mutations in cancers and normal cells correlates with the age of the individual. In addition to these two known signatures, we revealed a Signature C characterized by C>T transitions at GpCpN trinucleotides, especially GpCpC sites. In particular, Signature C was found to be the major contributor to somatic mutations detected in more than 30 normal samples (Fig 3C).

### Epigenetic regulation may influence the proportion of mutational signatures in normal tissues

To identify genetic factors contributing to these mutational processes, we further performed whole-exome association analysis with age as a covariate and the frequency of Signature A, B and C as the quantitative value respectively, in our 40 unrelated normal samples (Methods). In total, 21 SNPs located in 18 protein coding genes were shown to correlate with the proportions of Signature B (p < 1×10^−10^, permutation test; Fig 3D and Table 1). Among these, 2 genes contained SET domain (enrichment P=0.031), which is an important sequence feature of putative methyl transferase involved in histone methylation. The 2 genes are *PRDM9* (PR Domain 9), a zinc finger protein catalyzes the trimethylation of histone H3 lysine 4 (H3K4me3) and *KMT2C*, a histone methyltransferase involve in leukemogenesis and developmental disorder [28-30]. This result indicates mutations in epigenetic regulators may influence the proportion of signature B in normal tissue. Moreover, two SNPs in *NOTCH2* are associated with the proportion of Signature B and C, respectively (Table 1). *NOTCH2* is a key member of notch signaling pathway, which is important in metazoan development and tissue renewal. Its inter-cellular domain can act as a transcription factor regulates cell proliferation through controlling the expression of cycling D1[31, 32]. Additionally, there is no SNPs were significant associated with Signature A. These results demonstrate genetic background may influence the mutational profile of each individual.

**Table 1.** Genes revealed by association analysis of signature frequency with whole-exome sequencing

| CHR | POS<sup>a</sup> | BETA<sup>b</sup> | STAT<sup>c</sup> | P | Adjust P<sup>d</sup> | GENE |
| --- | --- | --- | --- | --- | --- | --- |
| **Signature B** |  |  |  |  |  |  |
| 1 | 120,539,668 | 0.41 | 9.17 | 4.61×10<sup>-11</sup> | 6.55×10<sup>-8</sup> | *NOTCH2* |
| 5 | 23,527,777 | 0.64 | 9.21 | 4.13×10<sup>-11</sup> | 1.26×10<sup>-7</sup> | *PRDM9* |
| 7 | 151,945,204 | 0.55 | 9.39 | 2.51×10<sup>-11</sup> | 8.48×10<sup>-8</sup> | *KMT2C* |
| 7 | 151,962,309 | 0.43 | 9.60 | 1.40×10<sup>-11</sup> | 6.55×10<sup>-8</sup> | *KMT2C* |
| 9 | 33,386,243 | -0.81 | -10.36 | 1.75×10<sup>-12</sup> | 1.77×10<sup>-8</sup> | *AQP7* |
| 9 | 33,798,543 | 0.67 | 10.47 | 1.28×10<sup>-12</sup> | 1.77×10<sup>-8</sup> | *PRSS3* |
| 12 | 53,865,349 | 0.63 | 9.07 | 6.21×10<sup>-11</sup> | 1.40×10<sup>-7</sup> | *PCBP2* |
| 12 | 111,885,367 | 0.64 | 9.31 | 3.09×10<sup>-11</sup> | 9.90×10<sup>-8</sup> | *SH2B3* |
| 13 | 20,247,238 | -0.67 | -10.47 | 1.28×10<sup>-12</sup> | 1.77×10<sup>-8</sup> | *MPHOSPH8* |
| 13 | 25,671,429 | 1.11 | 9.09 | 5.79×10<sup>-11</sup> | 1.36×10<sup>-7</sup> | *PABPC3* |
| 16 | 33,410,688 | 0.55 | 9.39 | 2.51×10<sup>-11</sup> | 8.48×10<sup>-8</sup> | - |
| 17 | 21,319,682 | -0.67 | -10.93 | 3.87×10<sup>-13</sup> | 1.77×10<sup>-8</sup> | *KCNJ12* |
| 17 | 44,850,996 | 1.11 | 9.09 | 5.79×10<sup>-11</sup> | 1.36×10<sup>-7</sup> | *WNT3* |
| 17 | 45,214,558 | -0.67 | -10.47 | 1.28×10<sup>-12</sup> | 1.77×10<sup>-8</sup> | *CDC27* |
| 19 | 3,586,698 | 0.63 | 9.05 | 6.51×10<sup>-11</sup> | 1.41×10<sup>-7</sup> | *GIPC3* |
| 19 | 9,012,789 | 0.63 | 9.10 | 5.66×10<sup>-11</sup> | 1.36×10<sup>-7</sup> | *MUC16* |
| 19 | 17,734,390 | 1.11 | 9.09 | 5.79×10<sup>-11</sup> | 1.36×10<sup>-7</sup> | *UNC13A* |
| 20 | 26,094,525 | 1.11 | 9.09 | 5.79×10<sup>-11</sup> | 1.36×10<sup>-7</sup> | *NCOR1P1* |
| 20 | 29,625,935 | 0.55 | 9.39 | 2.51×10<sup>-11</sup> | 8.48×10<sup>-8</sup> | *FRG1BP* |
| 21 | 11,058,227 | 0.55 | 9.39 | 2.51×10<sup>-11</sup> | 8.48×10<sup>-8</sup> | *BAGE5* |
| 21 | 11,058,229 | 0.55 | 9.39 | 2.51×10<sup>-11</sup> | 8.48×10<sup>-8</sup> | *BAGE5* |
| **Signature C** |  |  |  |  |  |  |
| 1 | 120,539,687 | -0.42 | -9.04 | 6.59×10<sup>-11</sup> | 2.07×10<sup>-6</sup> | *NOTCH2* |

a. Human reference genome build GRCh37.
b. Regression coefficient.
c. Coefficient *t*-statistic.
d. Adjusted p-value was calculated by Benjamini & Hochberg (1995) step-up FDR control.

### Signature C was a development associated mutational type

Among all three mutational types identified above, Signature A and B have been reported to associate with age related process. However, such an age-related accumulation was not seen in our time point samples, i.e., sperm and blood, perhaps because of their liquid feature. Sequencing of liquid samples are likely to capture mutants in a large cell population with relatively high VAFs which may reflect their early occurrence during embryo development. In contrast, a solid tissue is maintained or regenerated by limited stem cells in a local region. Therefore, the late stage mutations mingled with early ones may be distinguished in sequencing data by VAF. As expected, mutations in muscle biopsy samples had significantly higher VAFs than those in blood and sperm samples (Fig 1C, P < 2.2×10^−16^). We then divided muscle mutations into high VAF group, which was more likely generated during embryo development, and low VAF group with the cutoff of VAF = 0.025. In high VAF group, a significantly high proportion of C>T at GpCpC sites, the Signature C, was observed (0.185 vs 0.06, S9 Fig), suggesting the association of this mutation type with the development.

### Hypoxia contributed to the occurrence of Signature C

We extracted sequencing datasets from The Cancer Genome Atlas (TCGA, Methods), including 3,827 samples from 22 types of tumor (S10 Fig) [2, 11, 33]. For the somatic mutation profiles, only a small proportion of TCGA tumor samples (4%, 153/3,827) exhibited high numbers of C>T transitions in the GpCpC context, of which 58 were clear cell renal cell carcinoma (CCRCC, S4 Table). Notably, they were distinguished from the rest of the 273 CCRCC samples in cluster analysis (Fig 4A) and their similarity to various types of normal tissue indicated that the same mutational processes occurred in both CCRCC and normal cells.

**Fig. 4.**
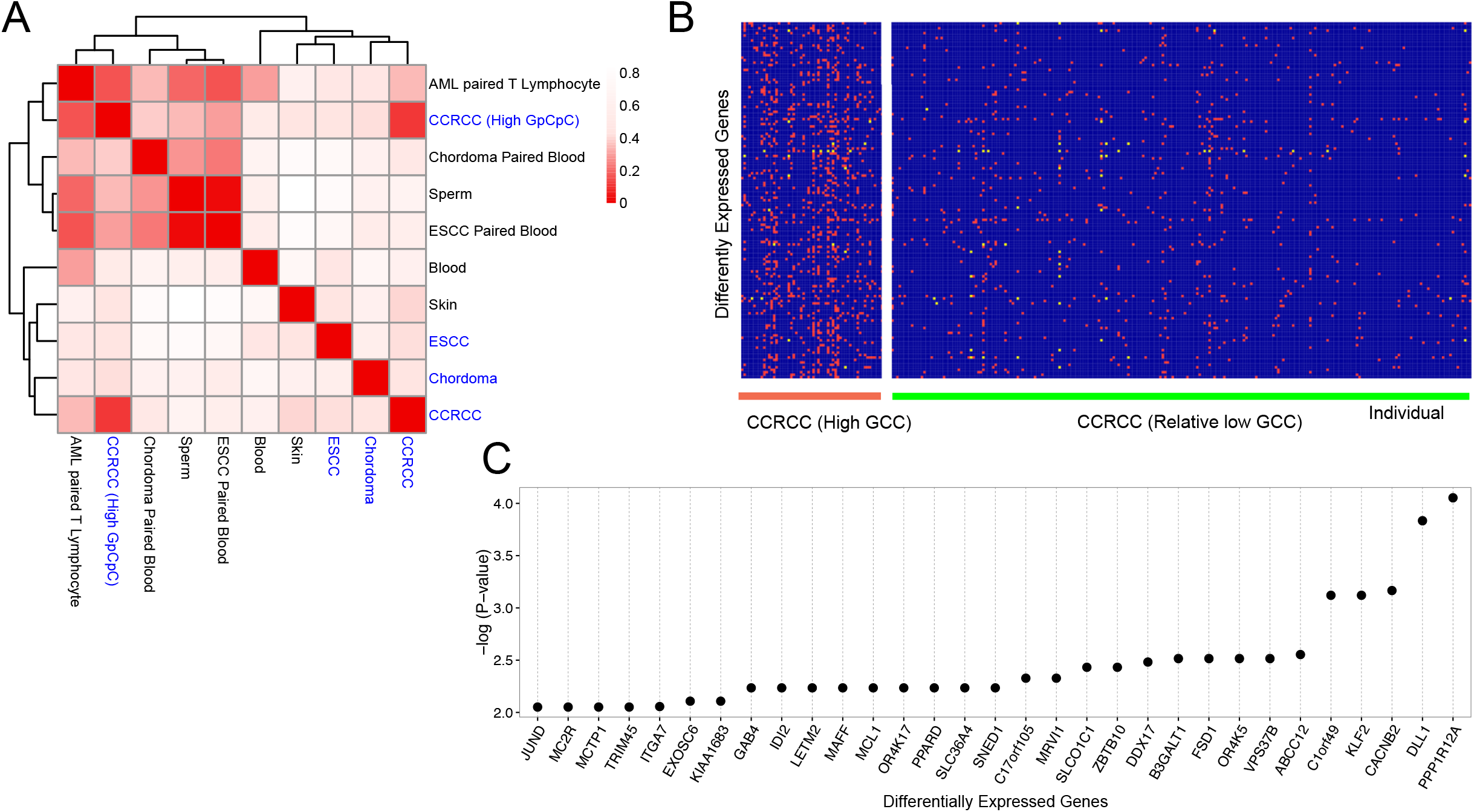
High GpCpC mutations in parts of CCRCC. (A) Correlation matrix of normal (black) and tumor (blue) cells. Among all tumors, a group of 58 CCRCC samples with high C>T transition at GpCpC were similar to normal cells regarding the mutational patterns. (B) Differentially expressed genes of CCRCC with high (left) and low GpCpC (right) after comparison with their adjacent normal tissues in TCGA. A total of 145 up- (red) and downregulated genes (yellow) were identified between the two groups. (C) Among the significantly differentially expressed genes (p < 0.01) in the high-GpCpC group, the top one *PPP1R12A* (p = 8.82×10^−5^) activates *HIF1A* by inhibiting HIF1AN-dependent suppression [34].

By comparing the expression profiles of high- (58 samples) and low-GpCpC groups (the remaining 237 samples) of CCRCC, we resolved 145 differentially expressed genes (p < 0.05, chi-squared test; Fig 4B, Methods). The most significant change in the high-GpCpC group was the increased transcription of PPP1R12A (protein phosphatase 1 regulatory subunit 12A), which activates hypoxia-inducible factor (HIF)-1α [34] (adjust-p = 8.82×10^−5^, Benjamini-Hochberg method, Fig 4C). Moreover, the Hippo pathway was significantly enriched (p = 3.21×10^−5^), which is associated with the transcriptional response to hypoxia [35-37]. Additionally, a slightly higher mutation rate of VHL (0.5 vs 0.43), whose product is involved in the ubiquitination and degradation of HIF proteins [38], was also observed in the high-GpCpC group (S5 Table). Taken together, these results suggest that increased activity of the HIF signaling pathway may contribute to the high proportion of C>T transitions at GpCpC in these CCRCC samples.

To test the roles of the HIF signaling pathway, we treated the human embryonic stem cell (hESC) line WA07 (WiCell Research Institute) with ML228, a direct activator of the HIF signaling pathway, through stabilizing and activating the nuclear translocation of HIF-1α [39] (Methods, Fig 5A). In the first stage, WA07 cells were divided into two groups with ∼1,000 cells in each. One group was treated with ML228 (0.125 nmol/ml) for 15 days, and the other was treated with mock as a control. For the second stage, 10 cells were randomly picked up from each group and expanded to ∼1,000 cells with or without ML228, before harvesting for exome sequencing with barcoding in library construction (Methods). As expected, a significantly high proportion of C>T transitions at GpCpN was observed in ML228-treated cells in comparison with the level in the control (0.17 vs. 0.07, p = 0.0091, chi-squared test; Fig 5B–C). According to their proportions, we divided all detected mutants into high- (VAF>0.05, mainly originating in the first stage) and low-allele-fraction mutations (VAF≤0.05, mainly generated from the expansion process in the second stage). For both types of mutation, higher accumulation of C>T transitions at GpCpN was observed in treated cells than in the control (0.12 vs. 0.06 in the high allele fraction and 0.20 vs. 0.08 in the low allele fraction; S11 Fig). These results demonstrate that activation of the HIF signaling pathway can lead to C>T transitions at GpCpN.

**Fig. 5.**
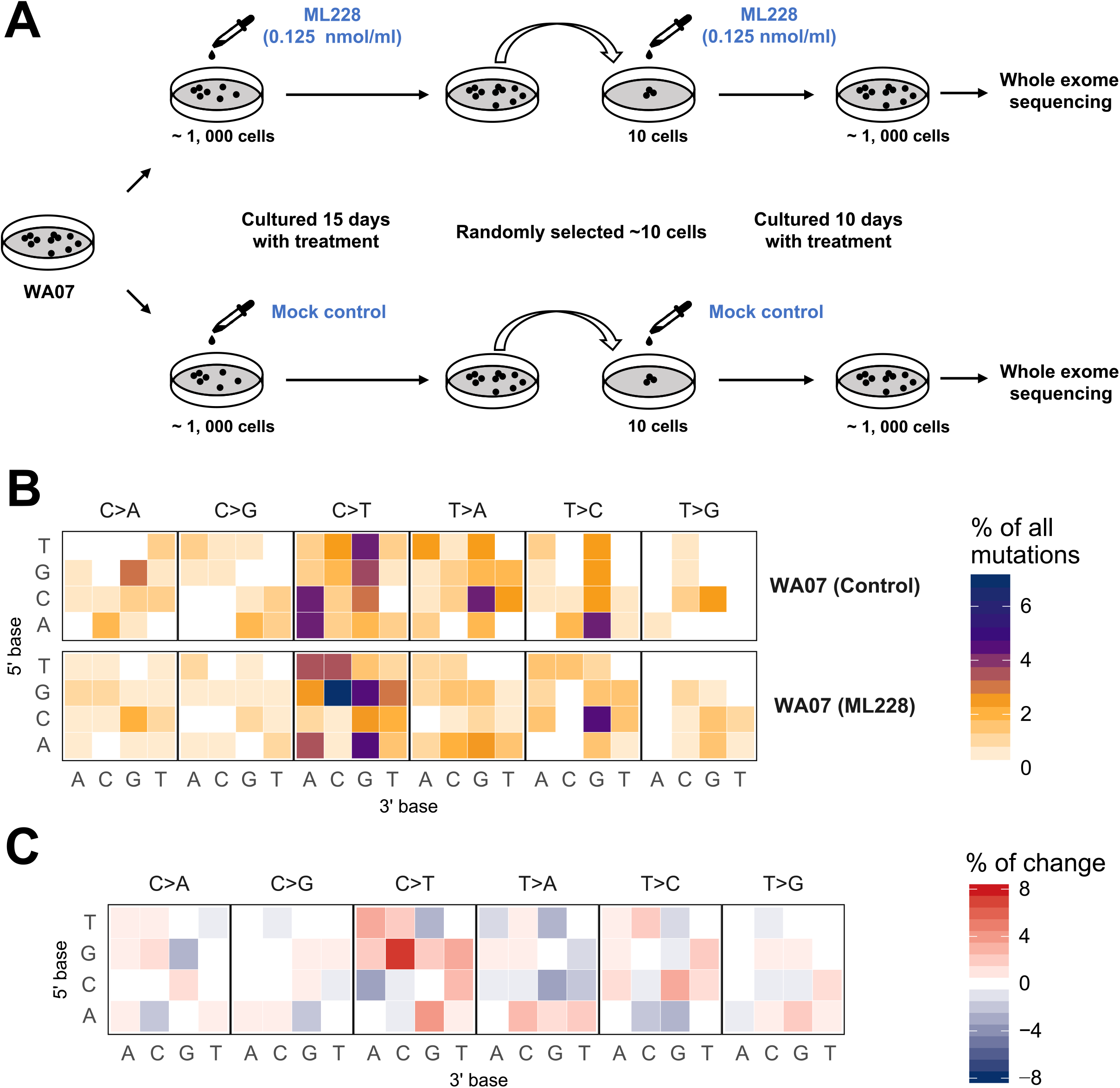
Activation of the HIF pathway by ML228 led to a high proportion of C>T transitions in the GpCpN context in hESC cells. (A) Two-stage treatment of WA07 cells with ML228 followed by exome sequencing with molecular barcoding. (B) A higher proportion of C>T transitions in the GpCpN context in ML228-treated cells. Note that the C>T transitions constituted the most significant difference in the GpCpC context of the ML228 group (0.07 vs. 0.01). (C) The fluctuation of 96 mutation types upon two-stage ML228 treatment. Among the 40 increased (red) and 26 decreased (blue) mutation types, C>T in the CpGpN context contributed to 14% of the total fluctuations, and half of this contribution was caused by the accumulation of C>T at GpGpC.

## Discussion

In this study, based on postzygotic mutation profiles in healthy individuals from five trio families, we discovered a signature characterized by C>T transitions in GpCpN trinucleotides as a major mutation type shared by blood and sperm cells. A solid evidence for this mutational pattern as a hallmark trait of normal tissues came from our observations in public or collaborative datasets eligible for analysis that such a signature was observed in all normal tissues but only very limited cancers. Furthermore, a portion of CCRCC with higher expression of HIF related pathways were featured by this mutation type, which lead to our speculation that the hypoxia status may trigger such mutations in healthy people. To prove this proposal, we designed an in vitro experiment using human embryo stem cell and we indeed observed accumulation of C>T transitions in GpCpN trinucleotides upon hypoxia induction.

Patterns of low VAF mutations in cell populations may be confounded by sequencing errors, and most sequencing artifacts are due to DNA damage during extraction and acoustic shearing [40, 41]. We used several strategies to assure the mutation authenticity. First, we largely reduced DNA damage before sequencing by introduction of repair mix (p < 0.05, S12 Fig, see Methods). We also used an optimized variant calling method to mask sequencing noise which allowed us to produce high-confidence calls of postzygotic mutations with VAF around 0.005 (See postzygotic mutation detection in Methods). Another challenge to identify postzygotic mutations is that under certain situations, it is difficult to distinguish inherited variants from postzygotic mutations due to inaccurate allele fractions in NGS sequencing. The theoretical values for allele fraction of inherited heterozygous variations should be 0.5, however measured values usually range from 0.2 to 0.6 due to unequal sequencing coverage of both alleles [42]. Actually, in our trio families, inherited mutations may have allele fractions even around 0.1 to 0.3 in the sequencing data from siblings (S13 Fig). Therefore, in addition to the filtering algorithms [43], we applied trio-based sequencing to preclude inherited mutations, and in this way common sequencing errors in multiple individuals can also be removed at the mean time. Validation of called variants with multiple methods ensured the confidence of our observation (S3 Table). Most importantly, in human embryo stem cells, we successfully generated our newly identified mutational signature by introduction of mutagenesis (Fig 5), which greatly supported the reliability of our findings.

To trace the mutagenesis process responsible for the signature, tumor samples in TCGA database provide us an interesting clue that this signature specific to normal tissues can be only found in a portion of CCRCC, known to be associated with activation of the HIF pathway [44]. Moreover, in recurrent glioblastoma, featured by extremely hypoxic conditions in tumor microenvironment [45], a recent study showed that C>T at GpCpC and GpCpT were enriched in their mutation spectrums (S2 Appendix and S14 Fig) [46]. Meanwhile, paired transcriptome analysis for the high-GpCpC group of CCRCC demonstrated that their HIF pathway was more active than that in the low-GpCpC group (Fig 4). The induction of GpCpC mutation by HIF signaling pathway was further validated in oligoclonal culture of hESCs. By directly activation of HIF1-α in oligoclonal hESCs using ML228, a previous well established assay [39], and a specially designed two-step cell culture experiments (Fig 5), we successfully observed the significant accumulation of C>T at GpCpN in mutational profiles.

In particular, we observed the same signatures for mutations in the blood and sperm tissues, indicating their common mutagenesis process. In fact, such C>T at GpCpC sites could also be seen in mutation profile of normal blood samples from a newborn baby study [47]. Notably, our samples carried high proportion of mutations in the range of 0.01∼0.05. This range of mutation fractions infer their occurrence within 20 cell divisions after fertilization. In view of HIF pathway being crucial in oxygen-sensing to mediate tissue adaptation to hypoxia, and hypoxic condition as a critical feature during embryonic development [48], we believe that the C>T transitions at GpCpN sites may occur during the embryonic development under the hypoxia status [49, 50]. In addition to our experimental validation with human stem cells, GpCpN mutations can also be observed in normal neurons, in which the accumulation of C>T at GpCpN mutations were significantly higher than those caused by deamination of methylated cytosines in NpCpG sites (Fig 3A, S2 Appendix, and S15 Fig, p=0.047, t-test) [26]. Since neuron cell division stops after the neuroepithelial cells have differentiated into proper neurons; most mutations should occur during cortical neurogenesis, which is complete around week 15 post-conception [51].

It is the two unique features in our sequencing strategy, the utilization of liquid samples sperm and blood, and the bulk sequencing without further separation or cloning, that allowed us to be able to capture those of relative common mutations in each tissue that occurred in the early stage of a cell lineage. In contrast, previous studies on postzygotic mutations mainly focusing on cancer somatic mutations, organoid mutations, or *de novo* mutations [1, 5, 52], most of which are private genomic changes in certain cell lineages across all the life span. In these mutation spectra, therefore, early events only account for a small proportion and possibly overwhelmed by all other mutations. This also explains why we observed the same signatures for mutations in the blood and sperm tissues, indicating their common mutagenesis process most likely during embryonic development.

Finally, integrating previously reported *de novo* mutations (S16 Fig), somatic mutations in CCRCC, and our results, we illustrate a mutational signature and corresponding active mutational processes during embryonic and post-parturition development, as well as in tumor development (Fig 6). Across the individuals’ lifespan, C>T transitions at NpCpG trinucleotides due to spontaneous deamination constantly occur after fertilization. In embryo development, the hypoxic environment triggers the occurrence and accumulation of C>T at GpCpN sites. After birth, T>C transition was generated in normal cells based on *de novo* mutations. In CCRCC, all types of mutation process were present. All of the aforementioned mutational processes can be observed in all tissue types. In future investigations, samples of multiple normal tissues from one individual should help to validate the molecular mechanism of hypoxia condition in mutation accumulation during embryonic development.

**Fig. 6.**
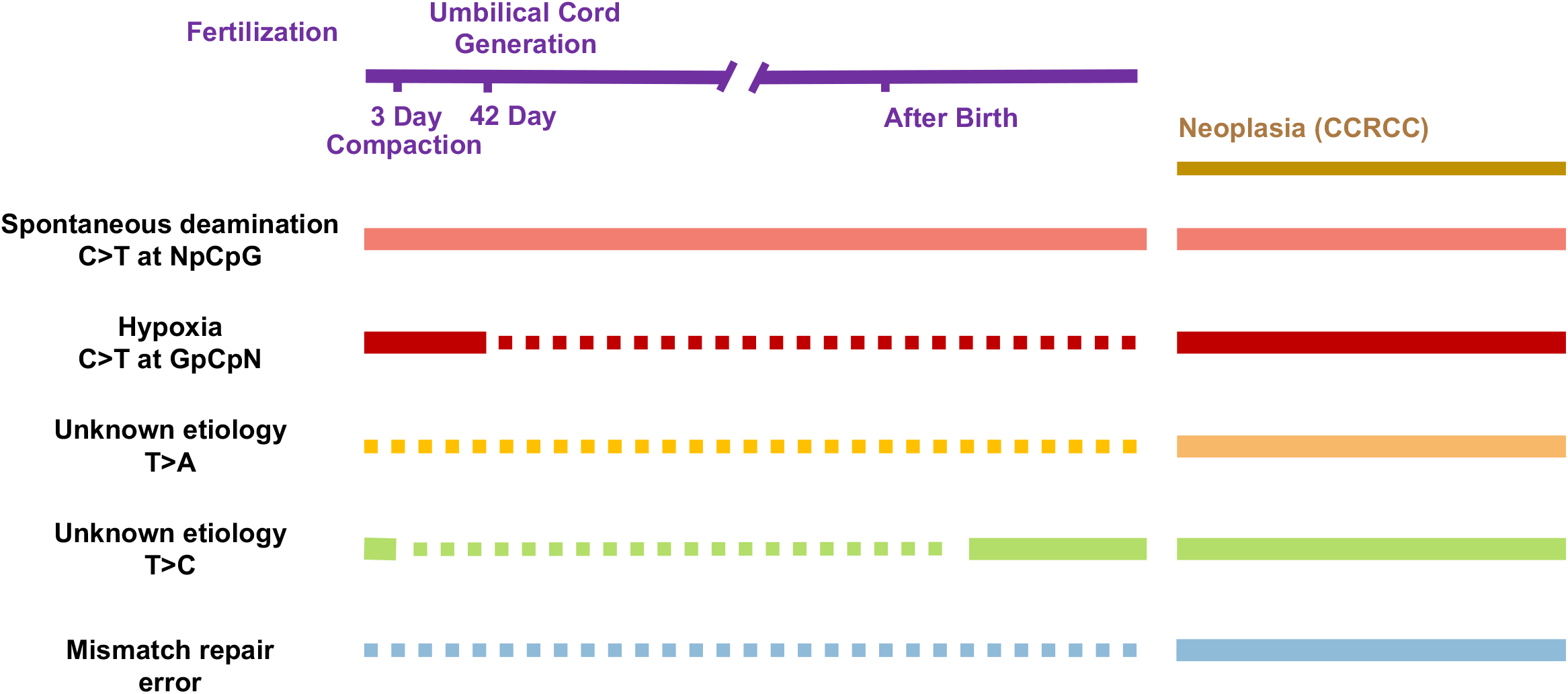
Proposed mutational processes (bottom) over the lifespan (top left) and in cancer of CCRCC (top right). After fertilization, the spontaneous deamination of methylated cytosine at NpCpG is the most common mutation type associated with age. In the early stage of embryonic development characterized by hypoxia, C>T transitions commonly occur at GpCpN sites. The enrichments of T>C transitions with unknown etiology are other mutational processes that occur during development. Regarding CCRCC development, the hypoxia-induced mutation process causes C>T transitions at GpCpN sites. Other mutational patterns in CCRCC include errors in mismatch repair and T>A transversions via unknown mechanisms. The dashed lines represent a lack of supporting evidence in a given stage.

## Materials and Methods

More detailed information is provided in S1 Appendix.

### Samples and whole exome sequencing

Individuals F0061, M0070, M0072, M0074, and their parents were enrolled at Wenzhou Medical University and samples from M0038 and his parents were collected at Beijing Institute of Genomics, Chinese Academy of Sciences (CAS). The five samples were 5-33 years of age and included four males and one female (S1 Table). All the details of whole exome sequencing analysis are summarized in S1 Appendix. This study was approved by the ethics committees of both Beijing Institute of Genomics, CAS (NO. 2016H006) and the Eye Hospital of Wenzhou Medical University (NO. KYK [2015] 2), and it was conducted in accordance with the principles of the Declaration of Helsinki principles. All the participants were healthy and provided written informed consent.

### Postzygotic mutation detection

Based on the error estimation model (detailed information is provided in S1 Appendix), postzygotic mutations in normal cells were detected by following several steps. Sequencing reads were aligned to the human reference genome build GRCh37 using the BWA algorithm [53] after the removal of adapter segments and the exclusion of reads with low Q-scores (S1 Appendix). Uniquely mapped reads with less than 3 mismatched bases were then processed using the error estimation model for all target regions, and variants with *P*^*m*^> *P*^*E*^ were selected out. Then, variants with more than 1% of reads supporting an alternative allele in either of the parents were removed to filter the inherited variants. In addition, due to the potential for misalignment, we only kept the variants included in the strict mask regions of the 1000 Genomes Project phase 1 [54].

### Mutational signature analysis

Mutational signatures were analyzed based on the guidelines of the Wellcome Trust Sanger Institute [11, 27]. The percentages of the 96 possible mutated trinucleotides in each sample, which were identified according to the six classes of base substitutions and 16 sequence contexts immediately 5’ and 3’ to the mutated base, were firstly calculated. The contexts of all mutations were extracted from the human reference genome build GRCh37. The mutational signatures in the selected samples were then estimated using the nonnegative matrix factorization (NMF) learning strategy. An appropriate number of mutational signatures was identified by calculating the reproducibility value and reconstruction error for all samples. Each mutational signature was finally displayed with the proportions of the 96 trinucleotides, and its contribution to each sample was estimated.

### Cell culture and molecular barcoded whole exome sequencing

The WA07 (WiCell Research Institute) cells were divided into two groups with ∼1,000 cells each and maintained in the human pluripotent stem cell chemical-defined medium (hPSC-CDM™, Baishou Biotechnology Co. LTD) according to the protocol. One group was treated with ML228 at 0.125 nmol/ml and the other group was treated with mock as the control. Both two groups were cultured for 15 day and cells received fresh medium with/without ML228 every other day. Then ∼10 cells were randomly selected from each group and cultured in the aforementioned medium with/without ML228 (0.125 nmol/ml), respectively. Cells received fresh medium with/without ML228 every five days. Molecular barcoded whole exome sequencing was performed on each group of cells after expanded to ∼1,000 cells. Genomic DNA of cultured expanded WA07 cells were extracted with a QIAamp DNA Mini Kit (Qiagen), per the manufacturer’s protocols. Partition barcoded libraries were then prepared based on the Chromium Exome Solution (10X Genomics) and the exome target regions was enriched by SureSelect Human All Exon V5 Kit (Agilent) according to the protocols. The target-enriched libraries with molecular barcoding were subsequently sequenced on a HiSeq 4000 (Illumina) with 150-bp paired-end reads.

### Mutation detection in hESCs

The exome sequencing data with molecular barcodes of WA07 cells was analyzed with the Long Ranger (10X Genomics). Then, the mutations which contained multiple molecular barcodes in the mismatched reads were kept. And to reduce the false positive rate, we removed the mutations which contained both two allele types in one molecular barcode in the site.

## Availability of data and materials

All sequencing data generated during the current study are available in the Genome Sequence Archive (http://gsa.big.ac.cn) with the accession number of CRA000071. Sources of the public tumor data used in this study are provided in the S1 Appendix. The unpublished data for ESCC, chordoma and AML samples is available upon request.

## Acknowledgments

The authors gratefully thank Dr. Ian M. Campbell and Dr. Pawel Stankiewicz from Baylor College of Medicine for kindly providing R source code for building cell-division model. The authors thank Dr. Xinyu Zhang from Yale University for critical reading and comments on the manuscript. The authors also thank Dr. Caixia Guo from Beijing Institute of Genomics, CAS for interpreting possible mechanisms underlying the occurrence of postzygotic mutations.

## Supporting Information

**S1 Appendix. Supplementary Methods**. The details of the methods in this study.

**S2 Appendix. Supplementary Notes**. The minor analysis and results in this study.

**S1 Fig. Pipeline for postzygotic mutation detection**.

**S2 Fig. Shared mutations among individuals and in time point samples of M0038**.

**S3 Fig. The distribution of variant allele fraction in each individual**.

**S4 Fig. QQ plot of the mutation fractions between simulation and observed in healthy individuals**.

**S5 Fig. Mutation type distribution in each individual**.

**S6 Fig. Mutation numbers of NpCpG and GpCpN among individuals of different ages**.

**S7 Fig. Replication asymmetry of each mutation type**.

**S8 Fig. Transcription asymmetry of each mutation type**.

**S9 Fig. Somatic mutation patterns in muscle**.

**S10 Fig. Somatic mutation patterns in TCGA samples**.

**S11 Fig. WA07 cells cultured with ML228 showed high fraction of C>T transitions at GpCpN context**.

**S12 Fig. The amounts of detected somatic mutations before and after repair mix treated**.

**S13 Fig. The distribution of inherited variant fraction**.

**S14 Fig. The mutational spectrum of recurrent glioblastoma**.

**S15 Fig. The mutational spectrum of normal neurons**.

**S16 Fig. The mutational spectrum of de novo mutations**.

**S17 Fig. The distribution of priori trinucleotide-specific sequencing error rate**.

**S18 Fig. The evaluation of mutagenic DNA damage induced sequencing error**.

**S19 Fig. Sequencing validation by duplicate reads**.

**S20 Fig. Possible bias among sequencing methods**.

**S21 Fig. The signature of hypermutated glioblastoma samples**.

**S22 Fig. The mutational signature of recurrent glioblastoma excluding hypermutated samples**.

**S23 Fig. No significant difference on DNA methylation was observed in CCRCC with high or low GpCpC mutations**.

**S24 Fig. The VHL mutation hotspots in high or low GpCpC group in CCRCC**.

**S1 Table. Sequenced healthy individuals**.

**S2 Table. Shared mutations in time point samples of M0038**.

**S3 Table. Validation list of mutations in the study**.

**S4 Table. Shared mutations among individuals**.

**S5 Table. Proportion of samples with C>T at GpCpC site as the major type in various cancers**.

**S6 Table. Mutations of VHL in CCRCC samples**.

**S7 Table. Features of somatic mutations compared with inherited mutations**.

**S8 Table. Somatic and inherited mutations of M0038**.

**S9 Table. Somatic and inherited mutations of F0061**.

**S10 Table. Somatic and inherited mutations of M0070**.

**S11 Table. Somatic and inherited mutations of M0072**.

**S12 Table. Somatic and inherited mutations of M0074**.

**S13 Table. Population frequency of mutations found in dbSNP (healthy individuals)**.

**S14 Table. Population frequency of mutations found in dbSNP (TCGA)**.

